# The role of stridulations during the mating of *Nicrophorus vespilloides*

**DOI:** 10.64898/2026.06.30.735517

**Authors:** Marie Guggenberger, Sophie Gerke, Taina Conrad

## Abstract

In many insect species, mating is coordinated through multimodal signaling, yet less obvious channels are often overlooked. In the burying beetle *Nicrophorus vespilloides*, chemical communication is well-documented, but the role of substrate-borne vibrational signals (stridulations) during courtship remains unknown. We investigated whether stridulation is essential for mating success through two sets of experiments. First, we found a positive correlation between the frequency of stridulations and both the number and duration of copulation events. Second, we employed a silencing experiment to test the necessity of these signals by silencing males, females, or both partners. We found no significant differences between silenced and control groups regarding the frequency or duration of physical contact and mounting events, suggesting that stridulation is not required for mate recognition or the initiation of courtship. However, the proportion of successful copulations relative to mounting events was significantly lower when females were silenced. These results suggest that while *N. vespilloides* relies on a redundant multimodal system that likely utilizes chemical cues to initiate mating, vibrational signals, particularly from the female, may play a critical role in facilitating successful copulation. This study provides the first evidence for the role of stridulation in the mating behavior *of N. vespilloides* and highlights the potential for female-mediated vibrational signaling in burying beetle courtship.

## Introduction

Communication occurs in various forms, such as visual, acoustic, vibrational, or chemical signals (Bradbury & Vehrencamp, 2011). Animals often rely on more than one communication channel at a time, a process known as multimodal signaling (Higham & Hebets, 2013; Lang, Conrad, Steiger, & Stökl, 2024; Mitoyen, Quigley, & Fusani, 2019; Partan & Marler, 1999).

Multimodal communication appears across a wide range of behaviors, allowing individuals to transmit richer or more robust information than a single modality alone could express, whether by reinforcing the same message or by carrying distinct information through each channel (Higham & Hebets, 2013). Courtship and mating represent one of the contexts in which multimodal communication is often observed (Lang et al., 2024; Ota, Gahr, & Soma, 2015; Stritih & Čokl, 2012; Virant-Doberlet, Stritih-Peljhan, Žunič-Kosi, & Polajnar, 2023). Birds, such as the blue-capped cordon-bleu (*Uraeginthus cyanocephalus*), perform courtship displays combining song with visual behaviours including bobbing and rapid step-dancing, the latter of which likely generates vibrations and/or non-vocal sounds (Ota et al., 2015). In the green stink bug *Nezara viridula* (Pentatomidae, Heteroptera), for instance, courtship relies on a sequence of signals, with long-range attraction mediated by male pheromones that lead to aggregation, followed by vibratory, visual, and chemical cues, through which both sexes initiate vibratory exchanges either spontaneously or in response to pheromones or visual stimuli (Zgonik & Čokl, 2014). Similar multimodal displays can be found in other insects (Eberhard & Picker, 2022; Hill, Mazzoni, Stritih-Peljhan, Virant-Doberlet, & Wessel, 2022; Stritih & Čokl, 2012).

However, in species where one mode has been intensively studied another modality is often overlooked. A prime example is the house cricket *Acheta domesticus* in which intensive study has focused on the auditory part of the signal (Stritih-Peljhan & Žunič-Kosi, 2024) only recently has it been discovered that they actually use a complex vibro-acoustic signal which might be the main condition dependent signal of the male.

In insects a strong focus in the past has been on chemical signals a critical role in a diverse array of behaviors including mating (Ayasse, Paxton, & Tengö, 2001; Steiger & Stökl, 2017; Stökl & Steiger, 2017; Wyatt, 2014). However, with the emerging field of biotremology (Hill et al., 2019; Hill et al., 2022) a new focus has been placed on vibratory signals in recent decades and more and more studies emerge showing that such signals often work in tandem with chemical signals (Conrad, 2022b; Hill, 2008; Lang et al., 2024; Virant-Doberlet & Cokl, 2004). Sophisticated stridulatory communication has for example been studied in the dung beetle *Aphodius ater*, where males produce complex, patterned vibrational songs upon encountering a female, with variation in song complexity suggesting a potential role in mate choice (Hirschberger, 2001).

One group which has been intensively studied regarding their chemical communication is the genus Nicrophorus (Silphidae). Studies were able to show that males release sex pheromones to attract females over long distances (Chemnitz, Bagrii, Ayasse, & Steiger, 2017; Chemnitz, Hoermann, Ayasse, & Steiger, 2020; Chemnitz, Jentschke, Ayasse, & Steiger, 2015; Haberer, Schmitt, Schreier, Eggert, & Müller, 2017; Steiger, Haberer, & Müller, 2011). It has also been found that the recognition of sex is dependent on their surface chemicals (Haberer, Steiger, & Müller, 2010; Keppner et al., 2017; Steiger, Whitlow, Peschke, & Müller, 2009). While the role of pheromones in mating is well documented, studies on the importance of stridulations during courtship and mating are lacking. Stridulations are produced by moving the plectrum on the ventral side of the elytra across the pars stridens, located on the fourth and fifth abdominal segments (Darwin, 1871).

Recent studies have shown that stridulations serve as a key communication mechanism during biparental care in Nicrophorus. For example, elytron clipping of *Nicrophorus americanus* for identification purposes influences one temporal and one spectral character of sound, which also decreases reproduction success (Hall, Howard, Smith, & Mason, 2015). It has also been shown that silencing of the parents and noise have a negative impact on offspring fitness (Conrad et al., 2024; Phillips, Chio, Hall, Hofstede, & Howard, 2020) and that stridulation increases significantly after hatching, which indicates, that this is an important part of the parental care in the *Nicrophorus vespilloides* (Conrad, 2022a). It is therefore plausible that stridulations also play a role in the mating of Nicrophorus and not only during brood care.

In this study we therefore set out to investigate whether, in addition to chemical signals, the substrate-borne communication in *N. vespilloides* is essential for their mating behaviour. We first recorded vibrations during mating and investigated any correlations with copulatory behaviour and then we investigated if the absence of stridulations during courtship influences the mating success, consisting of mounting duration, number of mounting events and number of successful copulations. We expected vibrational signals to play an important part in the mating behaviour of *Nicrophorus vespilloides* functioning as an additional means of information transfer for both partners.

## Materials and Methods

### Rearing and maintenance of beetles

Experimental beetles used were descendants of beetles collected from carrion-baited pitfall traps. *N. vespilloides* beetles were caught in a forest near Bayreuth, Germany (49°55’18.192’’N, 11°34’19.9488’’E). All beetles were maintained in temperature-controlled chambers at 20 °C on a 16:8 h light:dark cycle. Before the experiments, groups of up to 5 adults of the same sex and family of each species were kept in small plastic containers (10 × 10 cm and 6 cm high) filled with moist coconut coir. To ensure optimal outbreeding we used the program Kinshipper (Conrad, Bayreuth, Germany; kinshipper.com) to calculate optimal breeding pairs. Beetles were fed freshly cut larvae of both darkling beetles (*Zophobas morio*) or whole fly larvae (*Lucilia sericata*) ad libitum twice a week. At the time of our experiments, beetles were virgin and between 20 and 30 days of age.

### Behavioral recording of the animals

We recorded adult beetle pairs using HD TVI mini cameras (BSC TVI 2811, 2.8 - 12 mm, Eutin, Germany) with a frame rate of 25 and a resolution of 960 x 576. The cameras were connected to a recording device (LUPUS-Electronics GmbH, Landau, Germany). Acoustic signals were recorded using gooseneck microphones (t.bone GM 5212, Thomann, Burgebrach, Germany) connected to a computer via an audio interface (Tascam US-4x4 HR, Wiesbaden, Germany). This computer was equipped with a 32-bit sound card and the program Raven Ver. 1.4.1 (Center for Conservation Bioacoustics, Ithaca, NY, USA). The audio signals were digitized with a sampling rate of 44.1 kHz and a sampling format of 24-bit signed PCM.

The beetles were placed in pairs in transparent plastic boxes (17.5 x 11.5 x 7 cm), which will be referred to as sound boxes in the following. These consisted of a large (11.5 x 11.5 x 7 cm) and a small (6 x 11.5 x 7 cm) chamber separated by a metal grid.

The beetle pairs were located in the large chamber during the observation and the bottom of it was covered with coconut coir. The microphone was inserted into the smaller chamber. The sound boxes were placed in large, enclosed boxes with pyramid foam taped to the inner walls to reduce background noise. Video and audio recordings were then made of each pair of beetles under red light. During the recording, four sound boxes, each containing one pair, were in one sound proof box at the same time (Fig. 1).

**Fig. 1:**
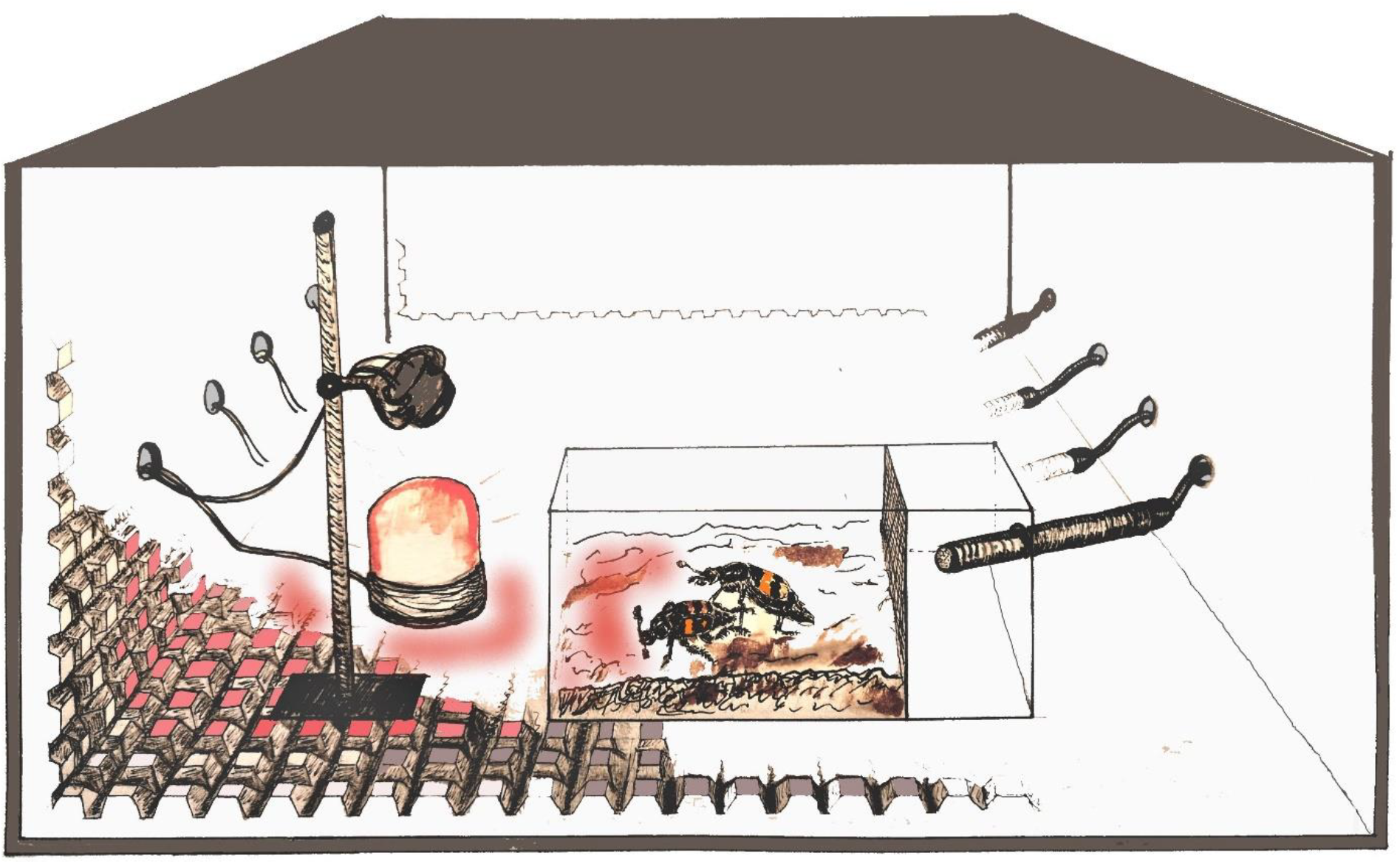
Pairs were recorded in transparent enclosures partitioned into a large observation chamber and a smaller compartment. The units were positioned within large foam-lined boxes to minimize background noise. HD-TVI cameras mounted above each enclosure captured video data, while microphones connected to a computer system recorded acoustic signals. Up to four pairs were monitored concurrently under red-light illumination.

### Mating behavior observations

In a first set of experiments, we observed mating and acoustic behavior in 24 couples of *N. vespilloides* during the first 90 minutes after pairing females and males. Videos and audio recordings were conducted between 15 November 2021 and 7 February 2022. We measured stridulation duration, copulation duration, and the number of copulation events for each couple.

### Experimental design for silenced couples

To analyze the effect of missing stridulations we silenced beetles and consequently observed their mating behavior in a second set of experiments. Mating pairs (n = 144) were chosen according to the program Kinshipper, their pronotum width documented with a stereo microscope equipped with a camera (Stemi 305, Zeiss, Berlin, Germany) and then assigned randomly to the silenced or control group. For silencing we used the same method as described in Conrad et al (2024). Beetles were anesthetized using CO_2_ and subsequently silenced by gluing a small (approx. 4mm) piece of parafilm (Bernis Inc., Neenah, Wisconsin, USA) onto the stridulatory organ using super glue (Super Glue Ultra Gel, Pattex ©, Henkel AG &Co KGaA, Düsseldorf, Germany). The control beetles were treated the same way but the parafilm was placed onto the lower part of the abdomen where it would not interfere with the stridulatory organ. After we attached the parafilm, beetles were kept anesthetized for approximately 10 more minutes to allow the glue to fully dry. Successful silencing was checked visually and audibly during handling throughout the experiment. Beetles were then recorded for two hours in the set-up mentioned above.

### Analysis of the video

We used the program Boris Ver. 8.19.3 (Department of Life Sciences and Systems Biology, University of Torino, Italy) to analyse the video recordings. Here, the number of mountings, the total and mean duration of mountings, the success of copulation attempts, and the length of physical contact of the beetle pairs were noted. A copulation attempt was included in the statistics as such and counted as mounting duration once the male attempted to assume the correct position. Copulation attempt was considered successful and counted as copulation when the male stayed in the correct position for a few moments and the beetle pair remained paused.

### Statistics

All analyses were done in R version 4.3.1 Graphs were produced using the R package ggplot, or the program Sigma Plot 16.0 (Systat Software, Chicago, IL, USA). Recordings of stridulations during mating without treatment were analyzed using spearman correlations. Stridulation time of 24 beetle couples was compared to both copulation time and copulation events.

For all response variables, we fitted models including treatment (silenced vs. control), male parent size, female parent size, carcass weight and temperature as fixed effects, as well as an interaction between male and female parent size:

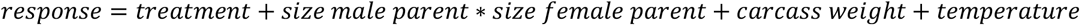

Gaussian models with an identity link function were used when model residuals were approximately normally distributed. For response variables that were strongly right-skewed, we fitted generalized linear models (GLMs) assuming a Gamma error distribution with a log link function. Count data (number of mounting events) were fit with a poisson distribution with a log link function.

Model residuals were inspected visually using standard diagnostic plots and by plotting residuals against fitted values and predictors. For GLMs, residual diagnostics were additionally assessed using the DHARMa package (v0.4.6; Hartig & Hartig, 2017). The contributions of different predictors to the variance in the data were tested via type II ANOVAs (linear models) and Likelihood Ratio Tests (GLMs) using the *Anova()* function (*car* package; Fox & Weisberg, 2019).

The evaluation was performed using the program RStudio Ver. 4.5.2 (Development Core Team, 2009).

Total mounting duration, mean mounting duration, number of mountings, copulation success, proportion of successful copulations and total time of physical contact were set as dependent variables. Treatment, parental size, and temperature and humidity during the observation period were fixed factors. If abnormalities were observed during the recordings, these data were excluded from the evaluation.

## Results

Under non-treatment conditions, mating *N. vespilloides* couples show a significant positive correlation between stridulation time and copulation time (n = 24, R = 0.7, p < 0.01, Fig. 2 A), as well as between stridulation time and the number of copulation events (n = 24, R = 0.71, p < 0.01 Fig. 2 B).

**Figure 2:**
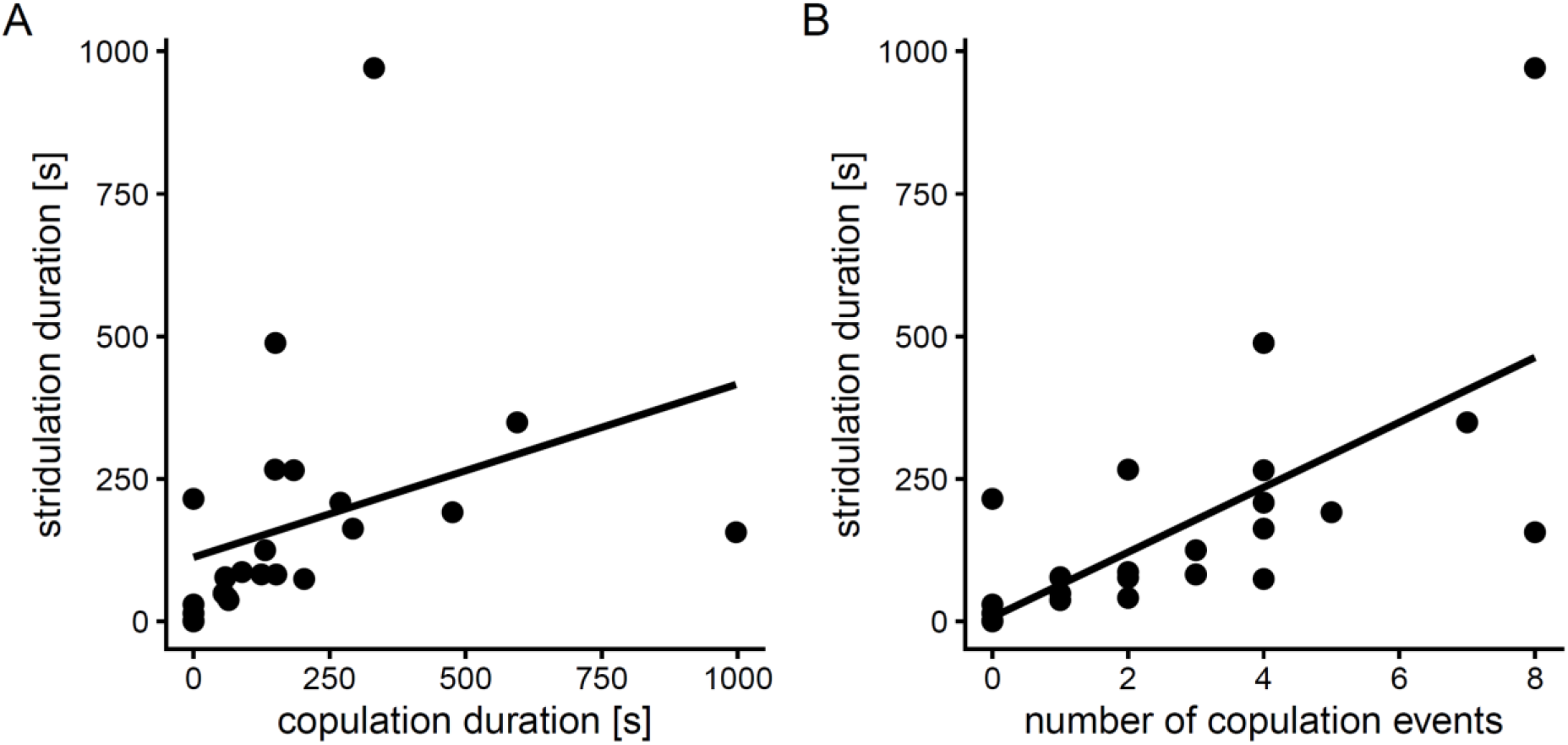
Stridulation time correlates with mating behavior. (A) Stridulation duration is positively correlated with copulation duration. (B) Stridulation duration is positively associated with the number of copulation events. Each point represents an individual couple (n = 24)

We found no significant difference between the treatment and the control groups in any of the three experiments when it comes to the time of physical touch (both silenced: GLM, ANOVA, F = 0.1669, p = 0.68529; male silenced: GLM, ANOVA, F = 0.0044, p = 0.9475; female silenced: GLM, ANOVA, F = 2.8138, p = 0.10385) (Fig. 3).

**Figure 3:**
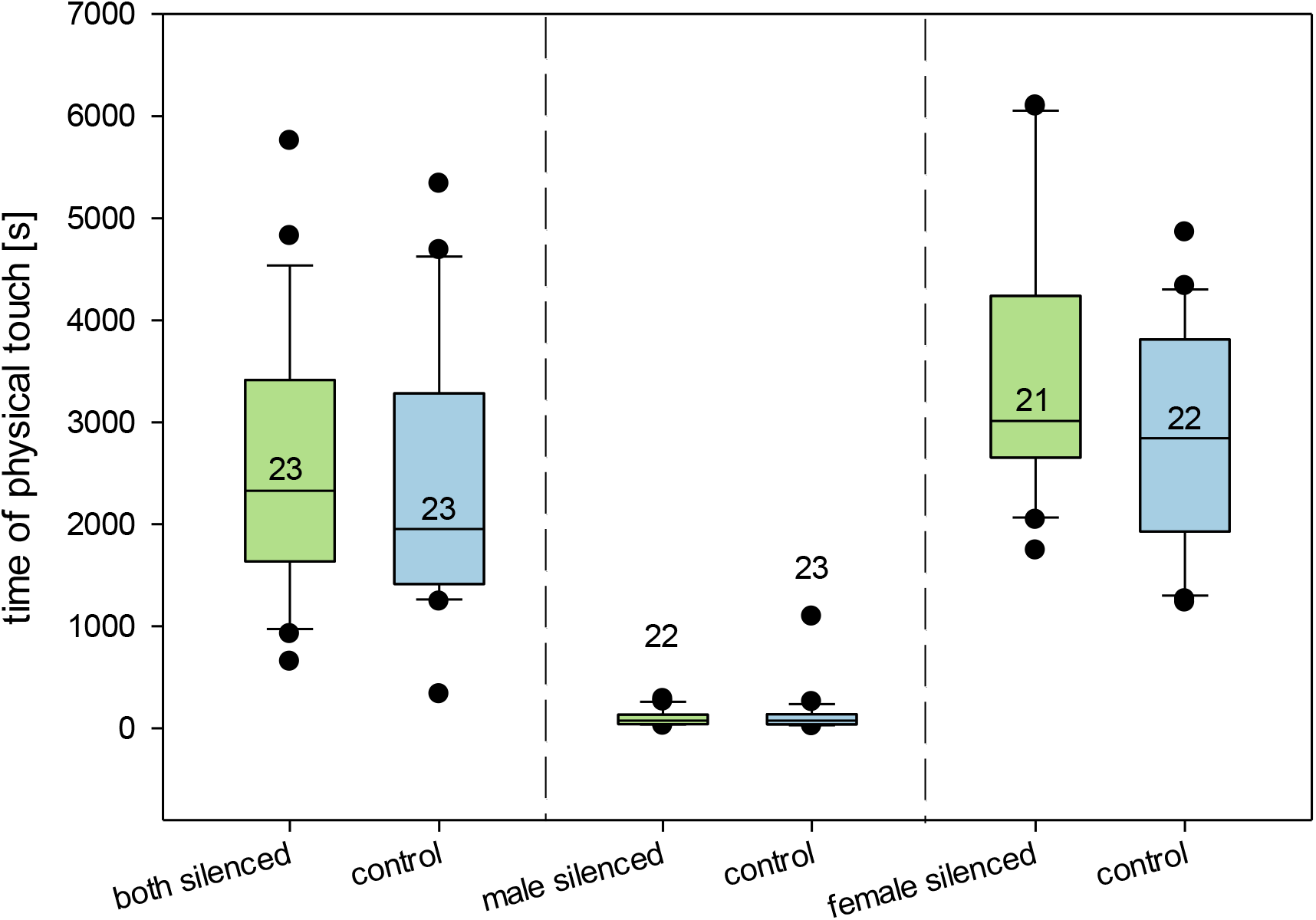
Time of physical touch of treatment group and control group of each experiment. The medians, quartiles and outliers (data points beyond 1.5×IQR from the quartiles) are shown. Whiskers represent the data range without outliers. Numbers represent the sample size for each group. Significant differences are marked by stars (GLM with subsequent Anova, *P < 0.05).

No effect could be shown for the average mounting duration either in both silenced (GLM; ANOVA, F = 3.4564, p = 0.07168), male silenced (GLM, ANOVA, F = 0.0063, p = 0.9373) or female silenced (GLM, ANOVA, F = 1.8504, p = 0.1839) (Fig. 4).

**Fig. 4:**
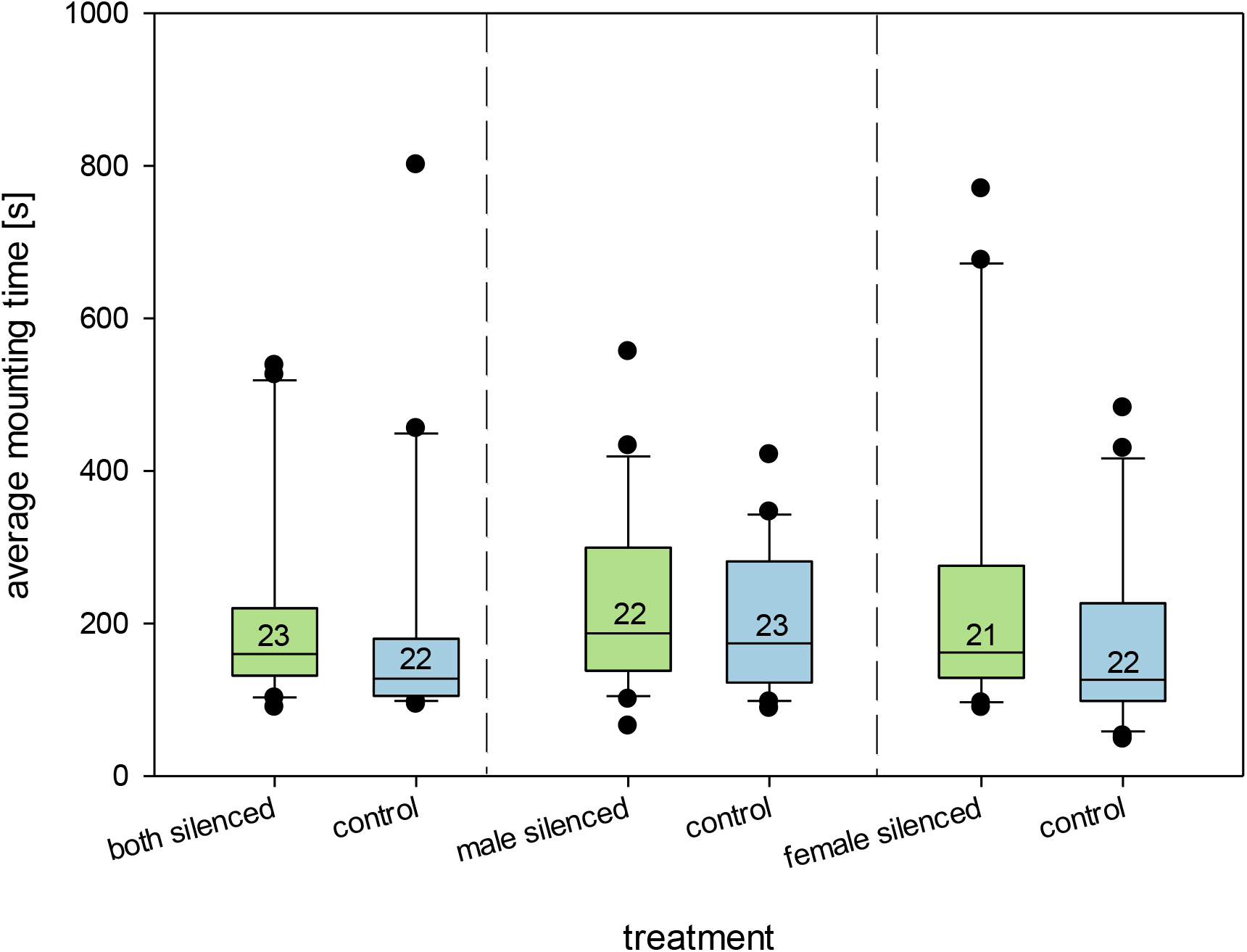
Average time of mounting of treatment group and control group of each experiment. The medians, quartiles and outliers (data points beyond 1.5×IQR from the quartiles) are shown. Whiskers represent the data range without outliers. Numbers represent the sample size for each group. Significant differences are marked by stars (GLM with subsequent Anova, *P < 0.05).

The comparison of the number of mounting events showed no significant differences for both silenced (GLM, ANOVA, Х^2^= 0.5393, p = 0.4627), male silenced (GLM; ANOVA, Х^2^= 0.4612, p = 0.4971) and female silenced (GLM, ANOVA, Х^2^ = 1.8206, p = 0.1772).

Looking at the success of copulations, there is a significant difference for the beetles in the treatment group of both silenced (GLM, ANOVA, Х^2^ = 4.1944, p = 0.0406) with the silenced group being less successful, than those in the control group. We didn’t find an effect in male silenced (GLM, ANOVA, Х^2^ = 0.2815, p = 0.5957) and in female silenced (GLM, ANOVA, Х^2^ = 2.9430, p = 0.0863), but the success of the beetles in these treatment groups was also lower (Fig. 5).

**Figure 5:**
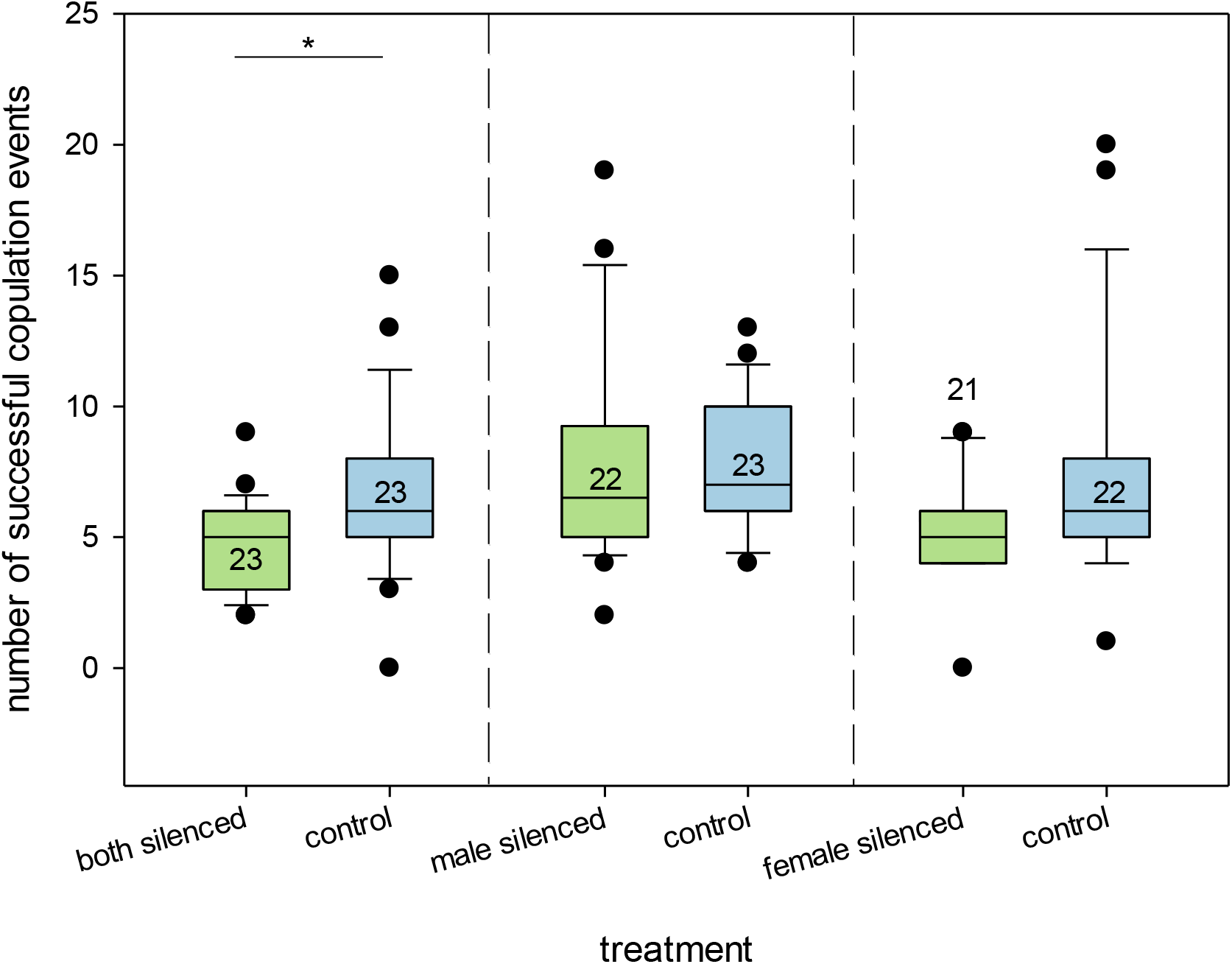
Number of successful copulation events of treatment group and control group of each experiment. The medians, quartiles and outliers (data points beyond 1.5×IQR from the quartiles) are shown. Whiskers represent the data range without outliers. Numbers represent the sample size for each group. Significant differences are marked by stars (GLM with subsequent Anova, *P < 0.05).

**Figure 6:**
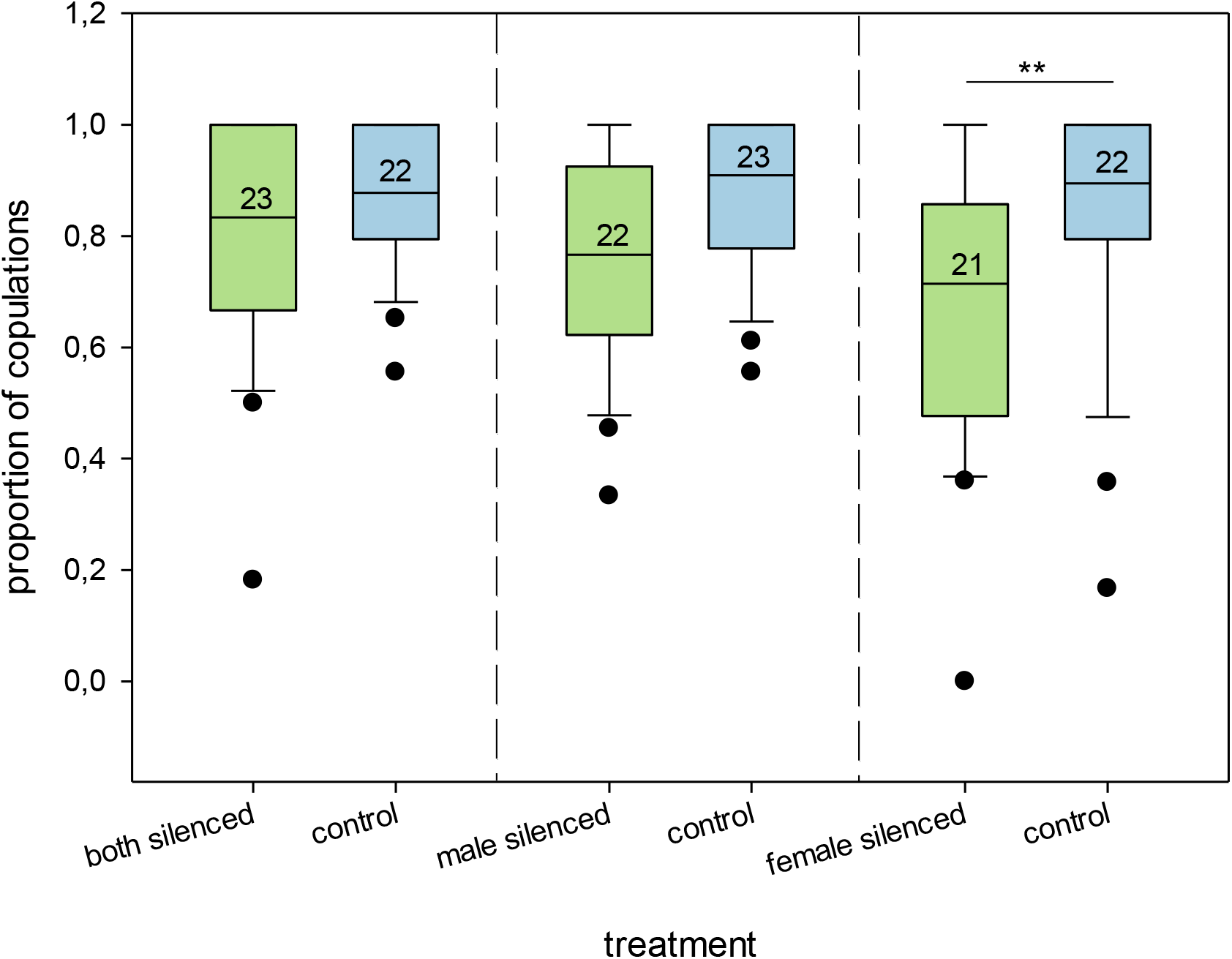
Proportion of copulation events to mounting events of treatment group and control group of each experiment. The medians, quartiles and outliers (data points beyond 1.5×IQR from the quartiles) are shown. Whiskers represent the data range without outliers. Numbers represent the sample size for each group. Significant differences are marked by stars (GLM with subsequent Anova, **P < 0.01).

For better comparability, the proportion of successful copulations to total mountings was calculated. A strong trend was found between the treatment group and the control group in both silenced (GLM, ANOVA, F =3.6334, p = 0.0648). In female silenced, the proportion of copulations is significantly lower in the treatment group (GLM, ANOVA, F = 8.3661, p = 0.007051). In male silenced, the proportion of copulations in the treatment group is not significantly different from the control (GLM, ANOVA, F = 2.0610, p = 0.1605) (Fig. 3).

## Discussion

Our results demonstrate a positive correlation between stridulation activity and successful copulation in *Nicrophorus vespilloides*. However, the lack of significant differences between silenced and control groups across most behavioral metrics suggests that the role of vibrational signaling in mating is more nuanced than we initially hypothesized.

Under control conditions, *Nicrophorus vespilloides* pairs that stridulated more frequently also showed a higher number of copulation events and longer total copulation durations. This positive correlation suggests that vibrational signaling is closely tied to the overall intensity of mating interactions. Similar patterns of signaling are observed across the genus *Nicrophorus*, indicating its widespread involvement in reproductive behavior. For instance, *N. nepalensis* males utilize specific stridulation frequencies to facilitate courtship, while in *N. americanus*, maintaining intact acoustic communication seems to be vital for achieving full reproductive success and optimal brood size (Hall et al., 2015; Hwang, Li, & Yang, 2024; Phillips et al., 2020). Interestingly, the silencing experiments revealed no significant differences in total physical contact, mounting duration, or the frequency of mounting events, regardless of whether one or both partners were silenced. The fact that females did not prevent mounting by silenced males, and those males did not terminate mounting when paired with silenced females, indicates that stridulation is not a prerequisite for mate recognition or the initiation of a copulation attempt. Instead, the vibrational signals may be a by-product of the mating interaction or a reinforcing signal rather than a trigger. However, the significant decrease in the proportion of successful copulations when both partners were silenced suggests that while stridulation is not necessary to start the process, it may be necessary to complete it.

We initially expected that stridulation would facilitate mate recognition, leading to more frequent encounters. In other insects, such as the genus Cenocorixa, stridulation is critical for mate recognition and female choice (Jansson, 1973), and in certain bark beetles, females use courtship songs to permit or reject copulation (Lindeman & Yack, 2015). In contrast, *N. vespilloides* appears to employ a less stringent mate-recognition strategy. This is consistent with the “acceptance threshold theory” (Reeve, 1989), which suggests that when the costs of mating are relatively low, individuals are less selective in their partner choice. It is further supported by the observed homosexual copulatory behavior in *N. vespilloides* (Engel, Männer, Ayasse, & Steiger, 2015), suggesting a low threshold for mating acceptance.

An alternative explanation for the resilience of mating behaviour in silenced beetles is the redundancy hypothesis (Hebets & Papaj, 2005) stating that in environments or settings where it is likely that one communication channel is disturbed it is advantageous to have a “backup” channel transmitting the same information content. Given the well-documented role of pheromones and surface hydrocarbons in Nicrophorus (Chemnitz et al., 2015; Keppner et al., 2017), it is likely that chemical cues provide sufficient information for mating to proceed in the absence of vibrational signals. The full necessity of stridulatory signals may only become apparent if chemical communication is simultaneously impaired.

The most striking result of this study is that the reduction in copulation success was specifically linked to the silencing of the female. This suggests a mechanism rooted not in female choice over a male signal, but in a signal produced by the female. It is plausible that females use stridulation to signal receptivity or to “permit” the male to copulate. In this scenario, the male can initiate mounting based on chemical cues, but the female must “answer” vibrationally to facilitate successful intromission. Alternatively, the absence of these signals could be interpreted by the male as an indicator of poor physical condition in the female, similar to male choice based on chemical cues in red turpentine beetles (Chen, Salcedo, & Sun, 2012) or pheromonal signals in the pink bollworm (Gonzalez-Karlsson et al., 2021). Because male mate choice is less frequently studied than female choice, the possibility that males use vibrational cues to assess female quality represents a significant area for future research.

Finally, it should be noted that our observations took place in relatively small sound boxes, which may have constrained the beetles’ interactions. To determine if silenced individuals are less efficient at locating each other over distance, future studies should utilize more spacious experimental setups.

In summary, this study demonstrates that while vibrational communication in *N. vespilloides* does not drive initial mate recognition or the initiation of mounting, it significantly influences the proportion of successful copulations—particularly when the female is silenced. These results suggest a multimodal mating system where vibrational signals, likely acting as a signal of female receptivity or quality, complement a robust chemical communication system. Future research should further investigate the interactions between these sensory modalities to fully clarify the complexities of burying beetle mating systems.

## Acknowledgements

We would like to thank Prof. Sandra Steiger for providing lab space and beetles. The research was supported by a grant of the German Research Foundation (DFG) to T.C (CO 2181/1-1 Project number: 424437154).

## CRediT

Marie Guggenberger (investigation, writing – original draft, formal analysis, visualization); Sophie Gerke (investigation, writing – original draft, formal analysis); Taina Conrad (conceptualization, funding acquisition, methodology, resources, supervision, formal analysis, visualization, writing – review & editing)

## Notes

### Competing Interest Statement

The authors have declared no competing interest.

